# Metagenomic Characterization of *Aedes aegypti* Virome in Kwale County, Kenya

**DOI:** 10.64898/2026.01.05.697663

**Authors:** Tabitha Wanjiru, Solomon Langat, Santos Yalwala, Gladys Kerich, Janet Ambale, Robert Haynes, John Eads, Fredrick Eyase

## Abstract

**Background:** *Aedes aegypti* is a primary vector of arboviruses, including dengue, chikungunya, yellow fever, and Zika, and is widespread along the Kenyan coast, a region with recurrent outbreaks. In addition to human-pathogenic arboviruses, *Ae. aegypti* harbors insect-specific viruses (ISVs) that replicate exclusively in arthropods and may influence mosquito physiology, immunity, and vector competence. However, data on ISVs in Kenyan *Ae. aegypti* populations are limited.

**Methods:** Twenty-nine mosquito pools containing 20 individual mosquitoes from Kwale County were homogenized, combined into a superpool, and subjected to total RNA extraction. Libraries were sequenced on an Illumina MiSeq platform. Initial analysis was done on CZ-ID platform. Reads were quality-filtered using PrinseqLite and assembled de novo with MEGAHIT. Phylogenetic analyses was performed with IQ-TREE using the Maximum Likelihood method.

**Results:** Metagenomic analysis revealed diverse ISVs in *Ae. aegypti*. Complete genomes of Fako virus and Tesano Aedes virus, and partial genomes of Aedes partiti-like virus, Cell fusing agent virus, and Formosus virus, were recovered. Genome lengths ranged from 1,149 to 10,146 nucleotides, with 91–99.5% identity to reference strains. Phylogenetic analysis placed the viruses within established ISV lineages, showing close evolutionary relationships with strains previously reported from Africa and other regions.

**Conclusions:** This study provides comprehensive characterization of ISVs in *Ae. aegypti* from Kwale County. The recovery of complete and near-complete genomes demonstrates the diversity and active circulation of ISVs, establishing a baseline for future studies on mosquito viromes, virus–mosquito interactions, and potential impacts on vector competence.

**Importance:** Insect-specific viruses (ISVs) are widespread in mosquitoes and replicate exclusively in arthropods, influencing mosquito physiology, immunity, and potentially vector competence for human-pathogenic arboviruses. Despite recurrent arboviral outbreaks along the Kenyan coast, the diversity and ecological roles of ISVs in *Aedes aegypti* remain poorly understood. This study provides a comprehensive metagenomic characterization of ISVs in *Ae. aegypti* from Kwale County, revealing complete genomes of Fako virus and Tesano Aedes virus, along with near-complete genomes of Aedes partiti-like virus, Cell fusing agent virus, and Formosus virus. These findings establish a baseline for mosquito virome composition in this region and underscore the active circulation of diverse ISVs in natural populations. Understanding these virus–mosquito interactions is critical for interpreting arbovirus ecology, assessing the potential impacts of ISVs on vector competence, and informing future vector surveillance and biological control strategies.

## Introduction

*Aedes aegypti* (Diptera: Culicidae) is a primary mosquito vector responsible for the transmission of several medically important arboviruses, including dengue virus (DENV), Zika virus (ZIKV), chikungunya virus (CHIKV), and yellow fever virus (YFV) (Omuoyo et al., 2023). These vector-borne pathogens represent significant public health challenges globally and in Kenya, where repeated outbreaks of dengue fever and other Aedes-transmitted diseases have been documented along the coastal and western regions (Courtney & Cranston, 2015). The ecology of *Ae. aegypti* is shaped by its adaptability to urban and peridomestic environments, high anthropophily, and ability to exploit diverse breeding habitats, making it a persistent and efficient arbovirus vector (Bhatt et al., 2013).

In addition to their role in transmitting vertebrate-infecting arboviruses, *Ae. aegypti* mosquitoes harbor a diverse assemblage of insect-specific viruses (ISVs), which are viruses that replicate exclusively in insects and are unable to infect vertebrate hosts (Carvalho & Long, 2021; Oguzie et al., 2022; Patterson et al., 2020). ISVs have been identified in multiple viral families, including *Flaviviridae*, *Togaviridae*, and others, reflecting extensive diversity within mosquito viromes (Amoa-Bosompem et al., 2020). The first ISV ever described was the cell fusing agent virus (CFAV), isolated from an *Ae. aegypti* cell culture, which does not replicate in vertebrate cells and is widely regarded as the prototype insect-specific virus (Bolling et al., 2015; Martin et al., 2019). Since then, advances in high-throughput sequencing and metagenomics have greatly expanded the catalogue of ISVs associated with mosquitoes (Langat, 2023).

Emerging evidence suggests that ISVs may influence mosquito biology and vector competence in complex ways. Some ISVs have been shown to modulate arbovirus replication, potentially through mechanisms such as superinfection exclusion or interactions with the mosquito immune system (Nasar et al., 2012; Vasilakis et al., 2013; Vasilakis & Tesh, 2015). Although the precise effects of many ISVs on arbovirus transmission are not fully resolved, experimental studies indicate that ISVs can either suppress or alter the replication dynamics of co-infecting arboviruses in mosquito hosts, with implications for disease transmission (Fish et al., 2017; Parry et al., 2021). These interactions highlight the importance of considering the broader viral ecology within mosquito populations when evaluating vector competence and arbovirus transmission risk.

Insect-specific viruses have increasingly attracted attention for their potential use in the development of novel vaccines. Because ISVs replicate exclusively in insect cells and are non-pathogenic to vertebrates, they provide a safe platform for the production of viral antigens and virus-like particles (VLPs) for immunization purposes (Erasmus et al., 2018; Hall-Mendelin et al., 2016; Hall et al., 2025). Recombinant ISVs expressing arboviral structural proteins have been shown to elicit protective immune responses in animal models without the risk of causing disease in humans (Nasar et al., 2015; Tan et al., 2023). Such approaches could serve as scalable and safe alternatives to conventional live-attenuated or inactivated vaccines, particularly for arboviruses such as dengue, Zika, and chikungunya that remain significant public health threats in endemic regions including Kenya.

In addition to their ecological roles, ISVs have significant potential in the development of ELISA-based diagnostic tools for arbovirus surveillance (Erasmus et al., 2015). Because ISV-based chimeric constructs can express structural antigens of vertebrate-infecting viruses while remaining replication-restricted in vertebrate cells, they provide a safe alternative antigen source for use in immunoassays. For example, chimeras based on the insect-restricted Eilat virus have been successfully used as high-quality antigen in IgM ELISA formats for chikungunya virus, demonstrating sensitivity and specificity comparable to traditional assays while allowing handling at lower biosafety levels and reducing reliance on live pathogenic virus preparations (Erasmus et al., 2015). This approach enhances the feasibility of developing ELISA diagnostics that can be deployed in resource-limited settings, improve assay safety, and reduce costs associated with antigen production. ISV-based antigens may thus strengthen serological surveillance frameworks for arboviruses by providing robust, safe, and scalable tools for early detection and outbreak monitoring

Despite growing interest in the ecological and evolutionary roles of ISVs, genomic data and detailed characterization of these viruses from *Ae. aegypti* populations in East Africa remain limited. Metagenomic studies in Kenya have identified a variety of ISVs, including flavivirus-like agents and iflaviruses, underscoring the presence of diverse viral taxa co-circulating with pathogenic arboviruses in local mosquito populations (Chiuya et al., 2021; Langat et al., 2021; Omuoyo et al., 2023). However, comprehensive genomic analyses and phylogenetic studies of Kenyan ISVs are still scarce, leaving gaps in our understanding of their diversity, evolutionary relationships, and potential functional interactions with arboviruses. By documenting the genomic diversity and evolutionary relationships of ISVs, studies such as ours not only enrich baseline virome resources but also offer avenues for exploiting ISVs in public health interventions. These findings have implications for arboviral risk assessment, vector surveillance programs, and the design of innovative vector control and disease mitigation strategies. In regions like coastal Kenya, where Ae. aegypti-driven arboviral outbreaks are recurrent, understanding the interplay between ISVs and pathogenic arboviruses could ultimately contribute to reducing disease burden and improving outbreak preparedness.

## Methodology

### Ethical Approval

Ethical approval was obtained from the Kenya Medical Research Institute (KEMRI) Scientific and Ethics Review Unit (SERU) under protocol number KEMRI/SERU/CVR/4702 and WRAIR# 3101. Permission to conduct the study was granted by the National Council for Science, Technology, and Innovation (NACOSTI).

### Study area

The study was conducted in Kwale County, (4.1730°S, 39.4520°E) which lies along the coastal region with a tropical climate, high temperatures (26–32°C), and seasonal rainfall that create favorable conditions for *Aedes aegypti* proliferation.

**Figure 1.**
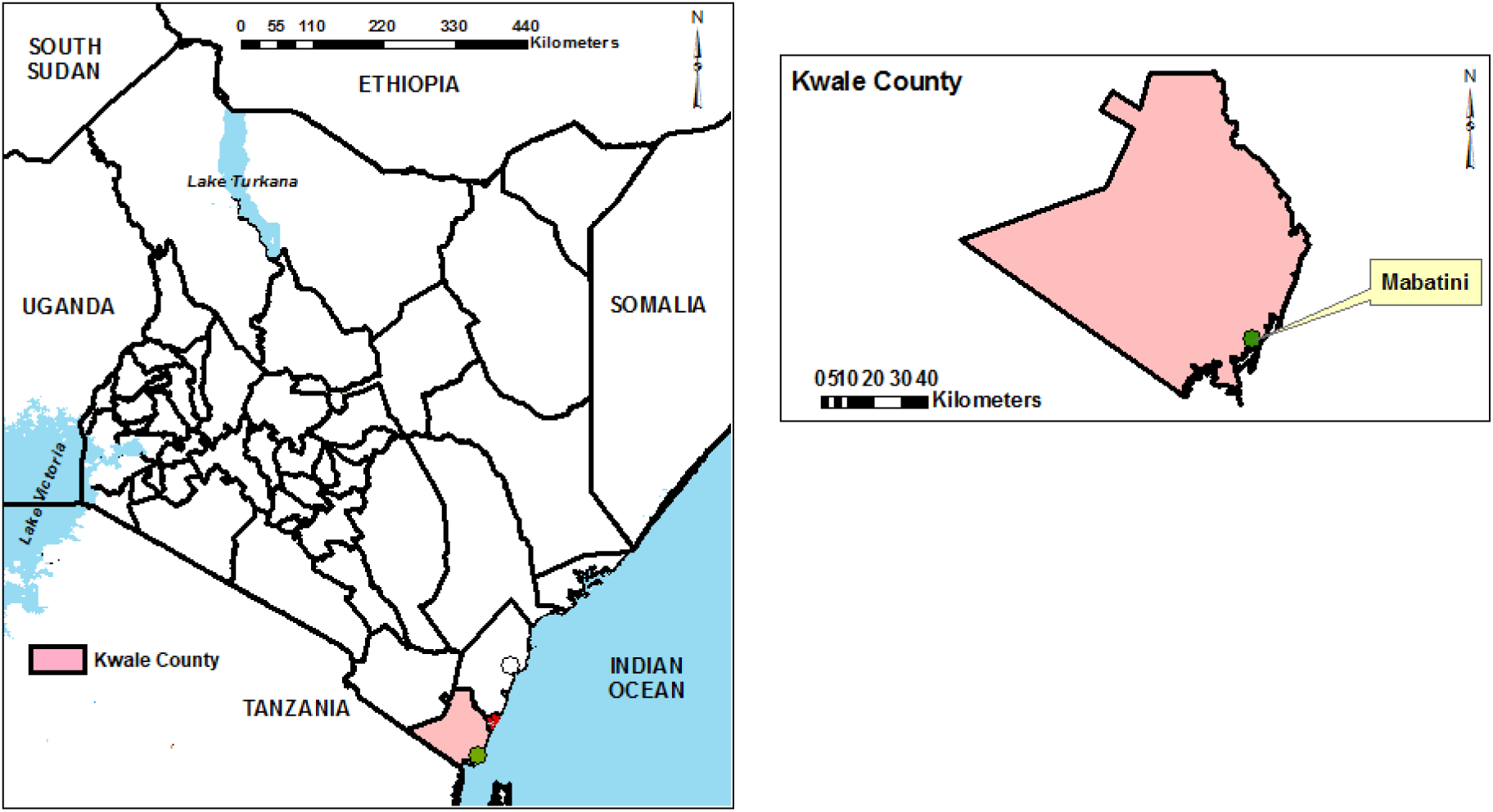
Map of the Kenyan coast showing mosquito sampling sites in Kwale County. Base maps, boundaries and shape files of Kenyan map and administrative boundaries of the Counties were derived from GADM data version 4.1 (https://gadm.org) and the maps were generated using ArcGIS Version 10.2.2 (http://desktop.arcgis.com/en/arcmap) advanced license) courtesy of Samuel Owaka.

### Entomological Investigation

Sampling of adult mosquitoes was conducted between 12^th^ March and 21^st^ June 2022. Mosquitoes were collected using CDC miniature light traps (Model 512, John Hock Co., Gainesville, Florida, USA). Traps baited with carbonated dry ice (CO2) were deployed overnight (6pm-6am) in favourable habitat, including dwelling quarters and animal sheds. The samples were linked to the sites by geo-coding using a GPS. The mosquitoes were immobilized by freezing at -20°C for 20mins and identified morphologically to species under a dissecting microscope using taxonomy keys, including Edwards (1941) (Road & Quaritch, 1936), Harbach (1988)(Harbach, 1988) and Jupp (1986) (Jupp, 1986). The identified mosquitoes were pooled in groups of 1 to 20 samples based on species, sex and collection site. Mosquitoes were consequently preserved in liquid nitrogen, and transported to the laboratory at the Kenya Medical Research Institute in Kisumu, where they were stored at-80°C until further processing.

### Mosquito sample preparation

A total of 580 mosquitoes collected from Kwale County were pooled into 29 pools, with each pool comprising 20 individual mosquitoes. The pools were homogenized using a Mini-Beadruptor-16 (BioSpec Products, Bartlesville, OK, USA) in 1,000 µL of homogenization medium, consisting of minimum essential medium supplemented with 15% fetal bovine serum (FBS) (Gibco, Life Technologies, Grand Island, NY, USA), 2% L-glutamine (Sigma-Aldrich), and 2% antibiotic–antimycotic solution (Gibco, Life Technologies), together with zirconium beads (2.0 mm diameter) for 40s. Homogenates were subsequently centrifuged at 10,000 rpm for 10 min at 4°C using a benchtop centrifuge (Eppendorf, USA). An aliquot of 50 µL of supernatant from each pool was combined to generate a single superpool for downstream metagenomic analysis.

### Library preparation and next generation sequencing

From the generated superpool, an aliquot of 140 µL of the supernatant was used for viral RNA extraction with the QIAamp Viral RNA Mini Kit (Qiagen, Hilden, Germany) and eluted in a single step with 60 µL of elution buffer, according to the manufacturer’s instructions. Paired-end sequencing libraries were prepared using the Illumina RNA Prep with Enrichment (L) Tagmentation (Illumina, USA) following the manufacturer’s recommended protocol. The final libraries were denatured with NaOH, diluted to a final concentration of 12 pM, and loaded onto an Illumina MiSeq platform. Sequencing was performed using the MiSeq Reagent Kit v3 (Illumina, USA) to generate 300-bp paired-end reads.

### Sequence analysis and virus identification

Initial analysis was performed using the CZ-ID platform, an integrated pipeline that offers quality control, de-hosting, duplicate removal, assembly, and viral identification capabilities. After this initial processing, the de-hosted sequence reads were retrieved for further analysis. To validate CZ-ID pipeline results, PrinseqLite v0.20.4 tool was used to filter low-quality reads and remove adapters on command line. De novo sequence assembly was conducted using MEGAHIT v1.2.9 (Li et al., 2015) where West Nile virus contigs were recovered and compared to CZ-ID pipeline results. Only contigs with an average depth of coverage of ≥10 and a length of ≥500 bp were retained for further analysis. These contigs were first compared against a local version of NCBI viral database using Diamond v2.0.4. To ensure specificity, putative viral contigs were further compared to the entire non-redundant protein database (nr), to exclude any non-viral contigs. A stringent e-value threshold of 1e-5 was employed throughout the homology searches to minimize false-positive hits.

### Phylogenetic analysis of the identified RNA viruses

To describe the identified viruses in an evolutionary context, publicly available viruses belonging to these different groups, and more specifically those closely related to the viral strains obtained in the current study were downloaded and used as reference sequences in the reconstruction of phylogenetic trees. Closely related gene sequences were retrieved from NCBI viral database and used as reference sequences in reconstructing the phylogenetic relationship of the viral sequences. The combined set of sequences were aligned using MUSCLE software with default parameters (max iterations = 16) embedded in Molecular Evolutionary Genetics Analysis v.7.0 (MEGA7) (Kumar et al., 2016) platform. The aligned sequences were edited using the Bioedit tool and maximum likelihood phylogenetic analysis carried out using IQ-TREE v1.6.12. The best model (GTR+G4 (General Time Reversible + Gamma)) and tree search was performed simultaneously based on 1000 bootstrap estimates and approximate likelihood ratio test (aLRT). Genome maps were generated and visualized using Python v3.9.

## Results

### Mosquito Collection and Sequencing Output

Adult *Aedes aegypti* mosquitoes collected from Kwale County, coastal Kenya, were processed for metagenomic analysis. Sequencing on the Illumina MiSeq platform generated 233,658 paired-end reads. After quality filtering, host read subtraction, and duplicate removal, 172,556 high-quality reads were retained for downstream virome analyses. Insect-specific viruses (ISVs) were detected exclusively in the Kwale County superpool. Superpools from Kilifi (Malindi) and Mombasa were also processed and sequenced, however, no sequencing reads were recovered, and consequently no insect-specific viruses or human-pathogenic arboviruses were detected.

### Metagenomic analysis

Metagenomic analysis revealed a diverse assemblage of insect-specific viruses (ISVs) associated with *Ae. aegypti* mosquitoes from the study area. Viral contigs were assigned to multiple taxonomic groups, representing both segmented and non-segmented RNA viruses. In total, five ISVs were identified, including Fako virus (FAKV), Tesano Aedes virus (TEAV), Aedes partiti-like virus (AePLV), Cell fusing agent virus (CFAV), and Formosus virus (FORV). Complete genome sequences were recovered for Fako virus and Tesano Aedes virus, while partial genome sequences were obtained for Aedes partiti-like virus, Cell fusing agent virus, and Formosus virus. Genomic features of the detected viruses, including genome length, GC content, closest reference sequences, and nucleotide identity, are summarized in Table 1.

**Table 1:**
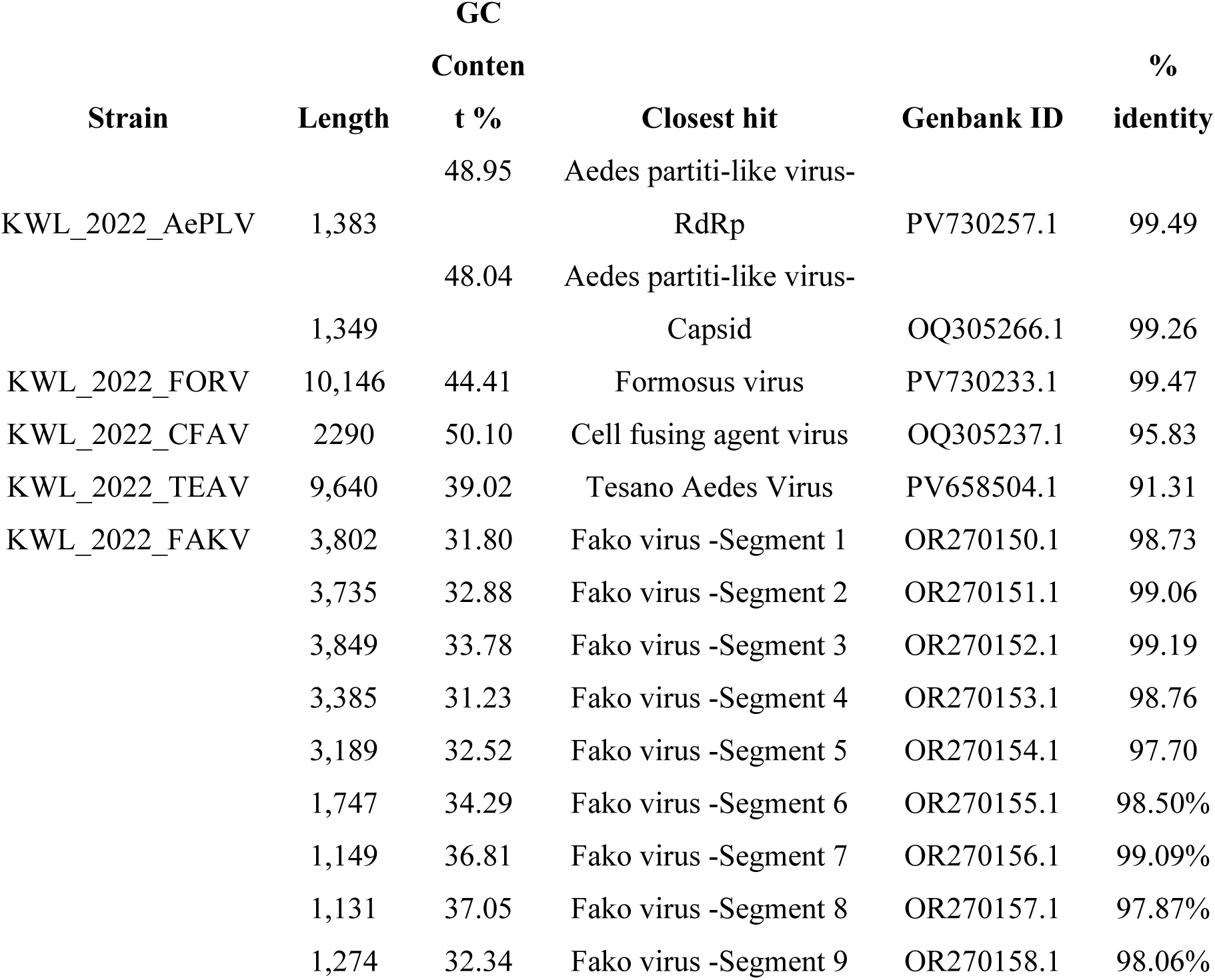
Viruses identified in this study based on metagenomic analysis. The identification was carried out using a homology search against reference databases, providing insights into the closest known virus.

### Phylogenetic analysis

Phylogenetic analyses based on the RNA-dependent RNA polymerase (RdRp) coding region were conducted to determine the evolutionary relationships of the detected ISVs. Maximum likelihood trees revealed that all identified viruses clustered within well-defined ISV clades, consistent with their taxonomic classifications. Aedes partiti-like virus was detected as two genomic segments corresponding to the RNA-dependent RNA polymerase (RdRp) and capsid regions. Both segments exhibited high nucleotide identity (>99%) to known AePLV reference sequences.

**Figure 2.**
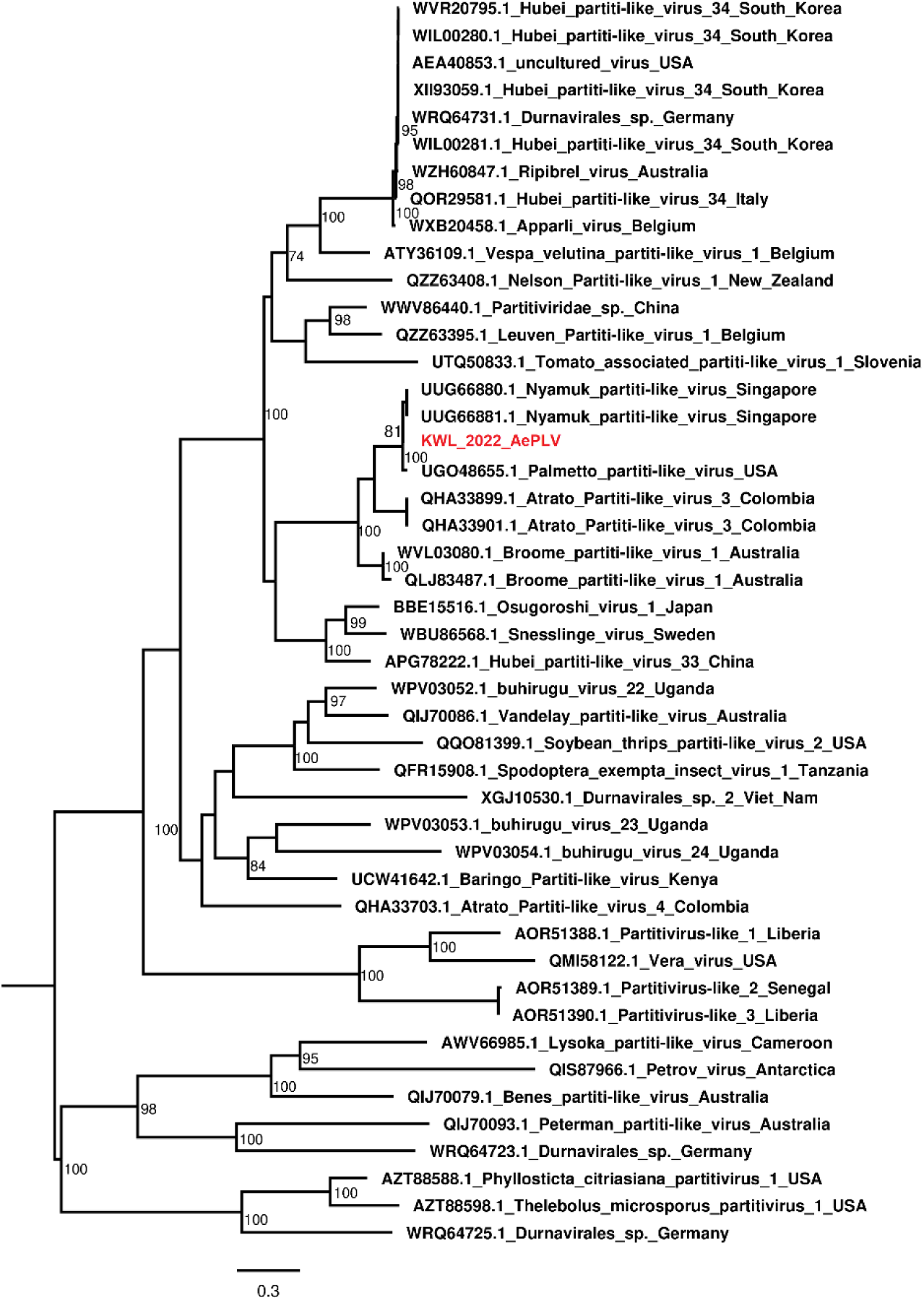
Maximum Likelihood phylogenetic tree of Aedes partiti-like virus (AePLV) inferred from nucleotide sequences, with branch support assessed using 1,000 bootstrap replicates.

### Genome organization of AePLV

The assembled genome of AePLV was determined to be bi-segmented, with lengths of 1,383 bp and 1,349 bp. Segment 1 encodes the RNA-dependent RNA polymerase (RdRp), whereas Segment 2 encodes the capsid and envelope-like proteins characteristic of insect-specific viruses. The genome maps reveal a linear organization for both segments, with conserved open reading frames (ORFs) and predicted untranslated regions (UTRs) at the 5’ and 3’ termini.

**Figure 3.**
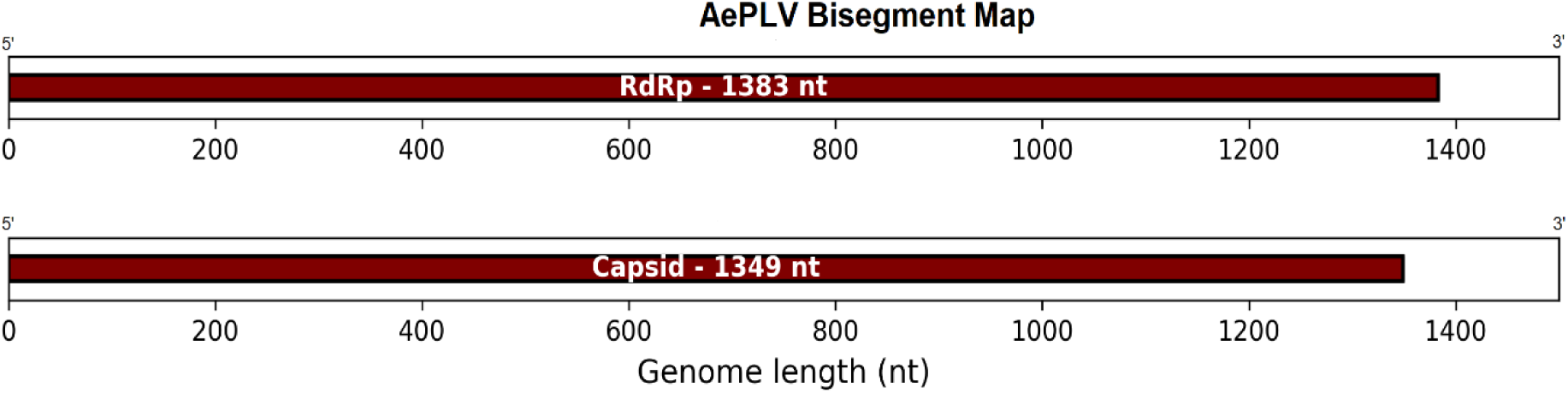
Genome map of AePLV showing two segments of varying lengths, each encoding a separate open reading frame.

Phylogenetic analysis of Formosus virus (FORV) showed that the recovered genome was 10,146 nucleotides in length with a GC content of 44.41%. The sequence exhibited high nucleotide similarity (99.47%) to the closest reference strain (PV730233.1) and clustered with a Nigerian strain, supported by a bootstrap value of 100.

**Figure 4.**
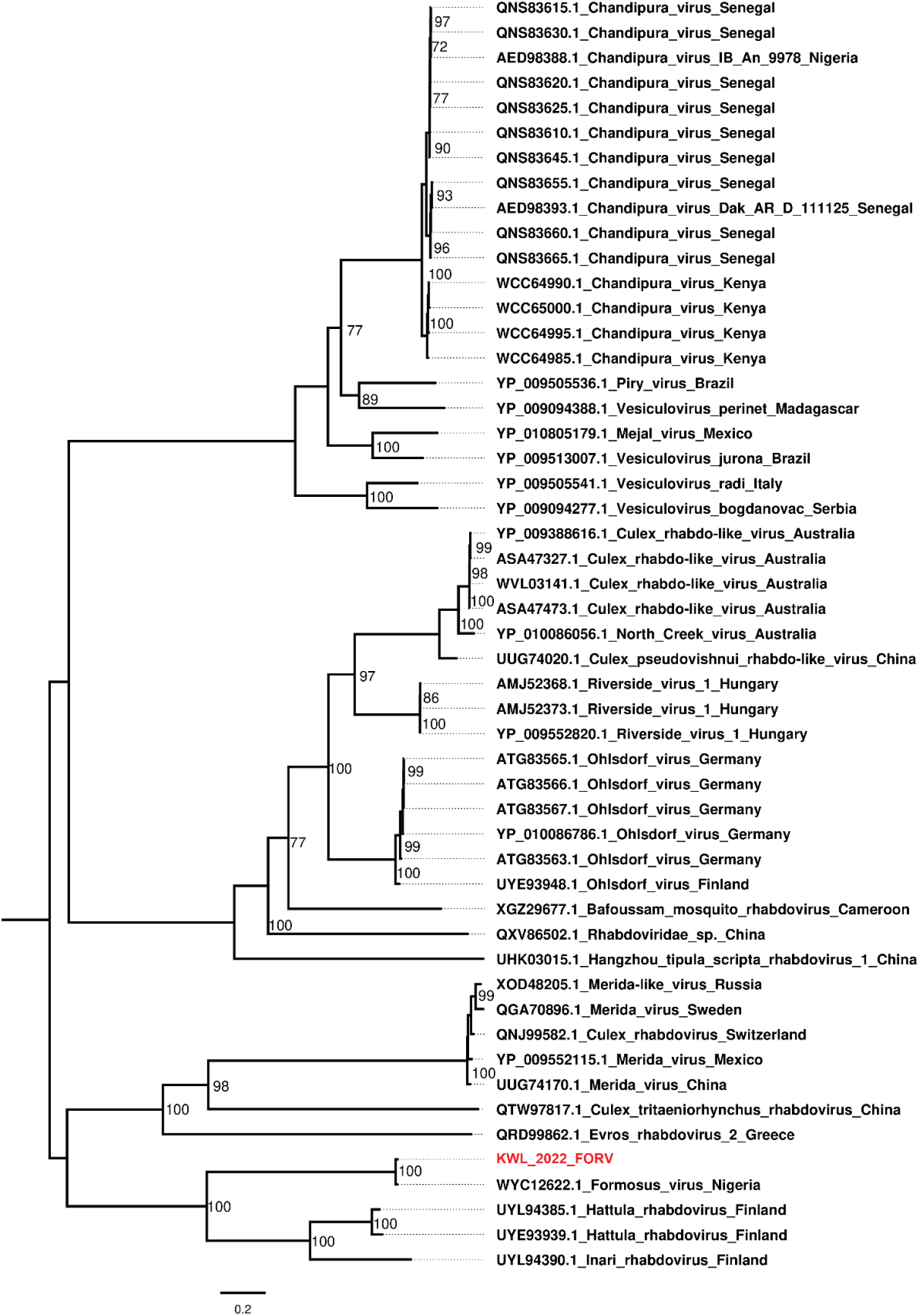
Maximum Likelihood phylogenetic tree of Formosus virus (FORV) inferred from nucleotide sequences, with branch support assessed using 1,000 bootstrap replicates.

The assembled genome of FORV consists of a single, continuous RNA segment of 10,146 bp. The genome encodes a complete RNA-dependent RNA polymerase (RdRp) and several structural proteins, including the capsid and putative envelope-associated proteins typical of insect-specific viruses. Analysis of the genome map revealed a linear organization with clearly defined open reading frames (ORFs) and untranslated regions (UTRs) at both the 5’ and 3’ ends.

**Figure 5.**
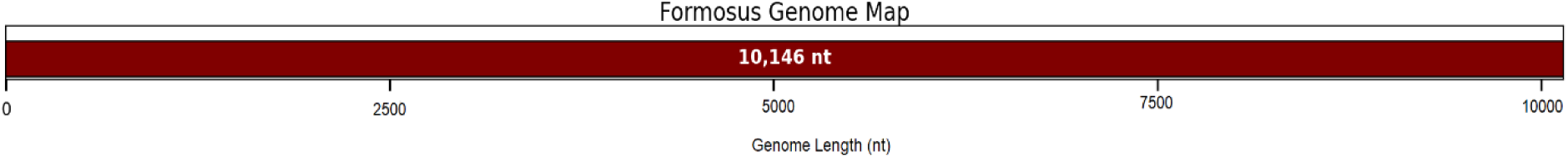
Genome map of Formosus virus (FORV) showing a single open reading frame in a genome of 10,146 nucleotides.

Phylogenetic analysis revealed that the Tesano Aedes virus genome clustered with a Kisumu-derived strain with a bootstrap value of 99. The genome showed nucleotide identity to the closest available reference indicating notable genetic divergence from previously reported strains.

**Figure 6.**
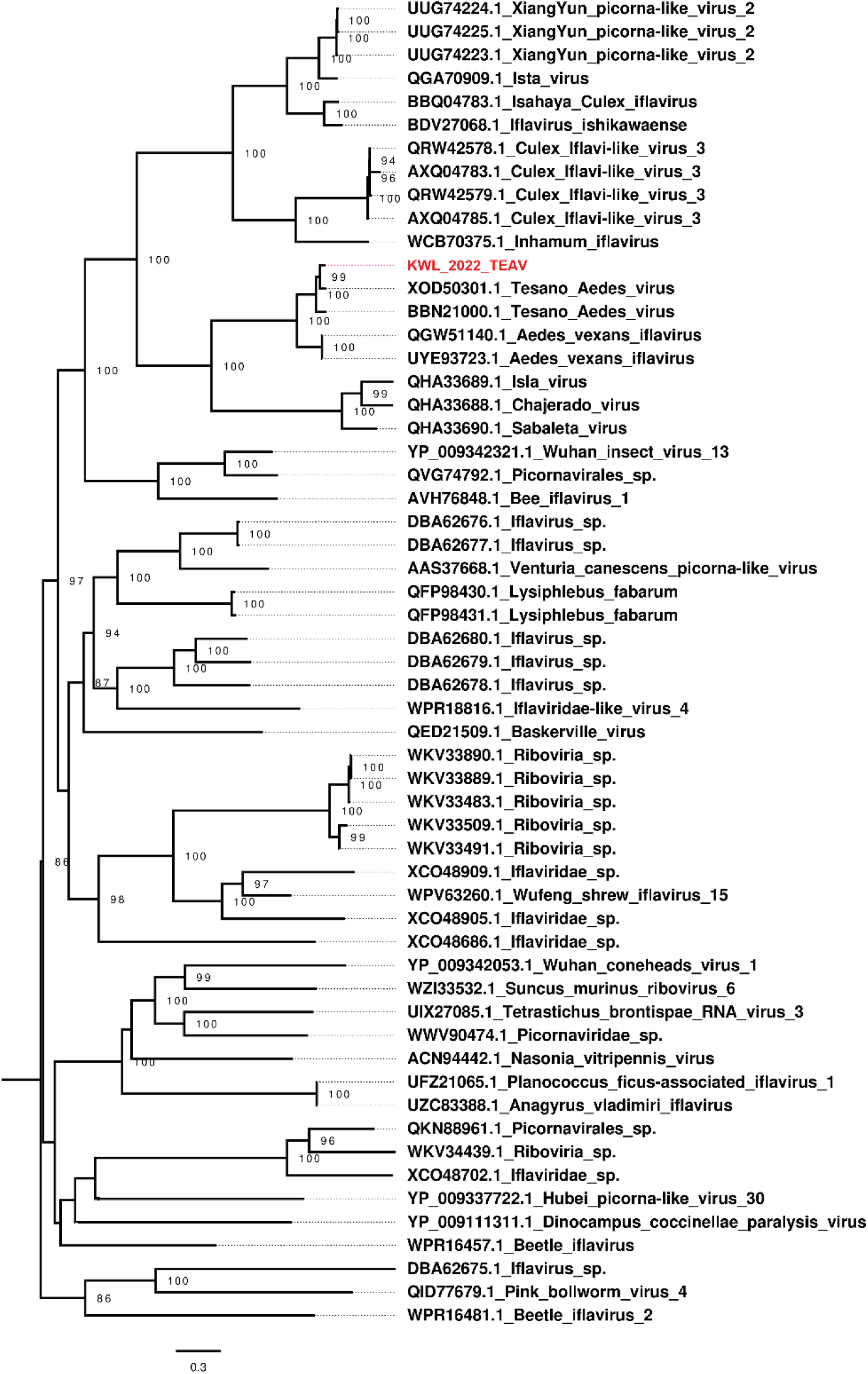
Maximum Likelihood phylogenetic tree of Tesano Aedes virus inferred from nucleotide sequences, with branch support assessed using 1,000 bootstrap replicates.

The Tesano Aedes virus genome recovered in this study was 9,640 nucleotides in length and contained a single open reading frame (ORF), characteristic of Iflaviruses, encoding a polyprotein that is post-translationally processed into structural and non-structural proteins.

**Figure 7.**
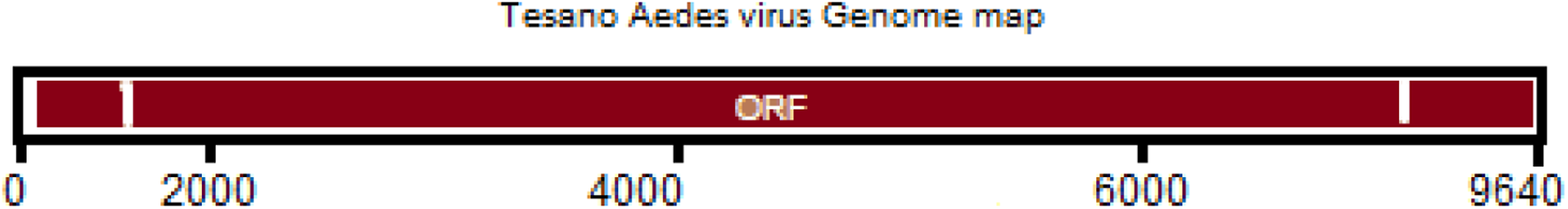
Genome map of Tesano Aedes virus (TEAV) depicting a single open reading frame in a genome of 9,640 nucleotides, characteristic of Iflaviruses.

Phylogenetic analysis of Fako virus revealed that the recovered genome clustered with a previously reported Kenyan strain, indicating close genetic relatedness. The branching was well supported in the phylogenetic tree, consistent with the virus circulating locally.

**Figure 8.**
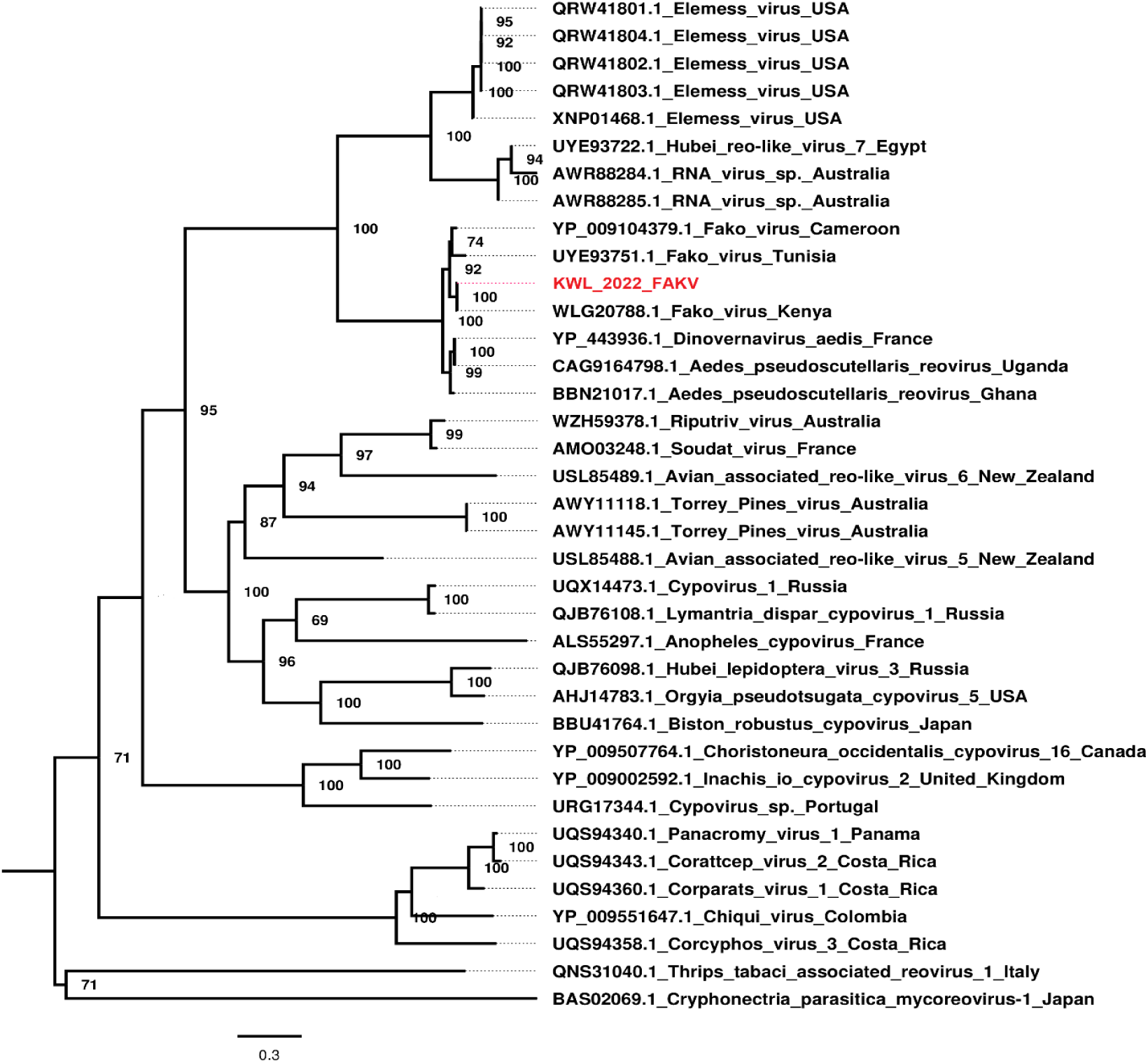
Maximum Likelihood phylogenetic tree of Fako virus (FAKV) inferred from RNA-dependent RNA polymerase (RdRp) gene sequences, with branch support assessed using 1,000 bootstrap replicates.

The Fako virus genome recovered in this study is segmented, comprising nine distinct segments of varying lengths. Each segment encodes a separate open reading frame, with the RNA-dependent RNA polymerase (RdRp) segment used for phylogenetic analysis. This segmented genome organization is consistent with previously described Fako virus strains and other related segmented viruses.

**Figure 9.**
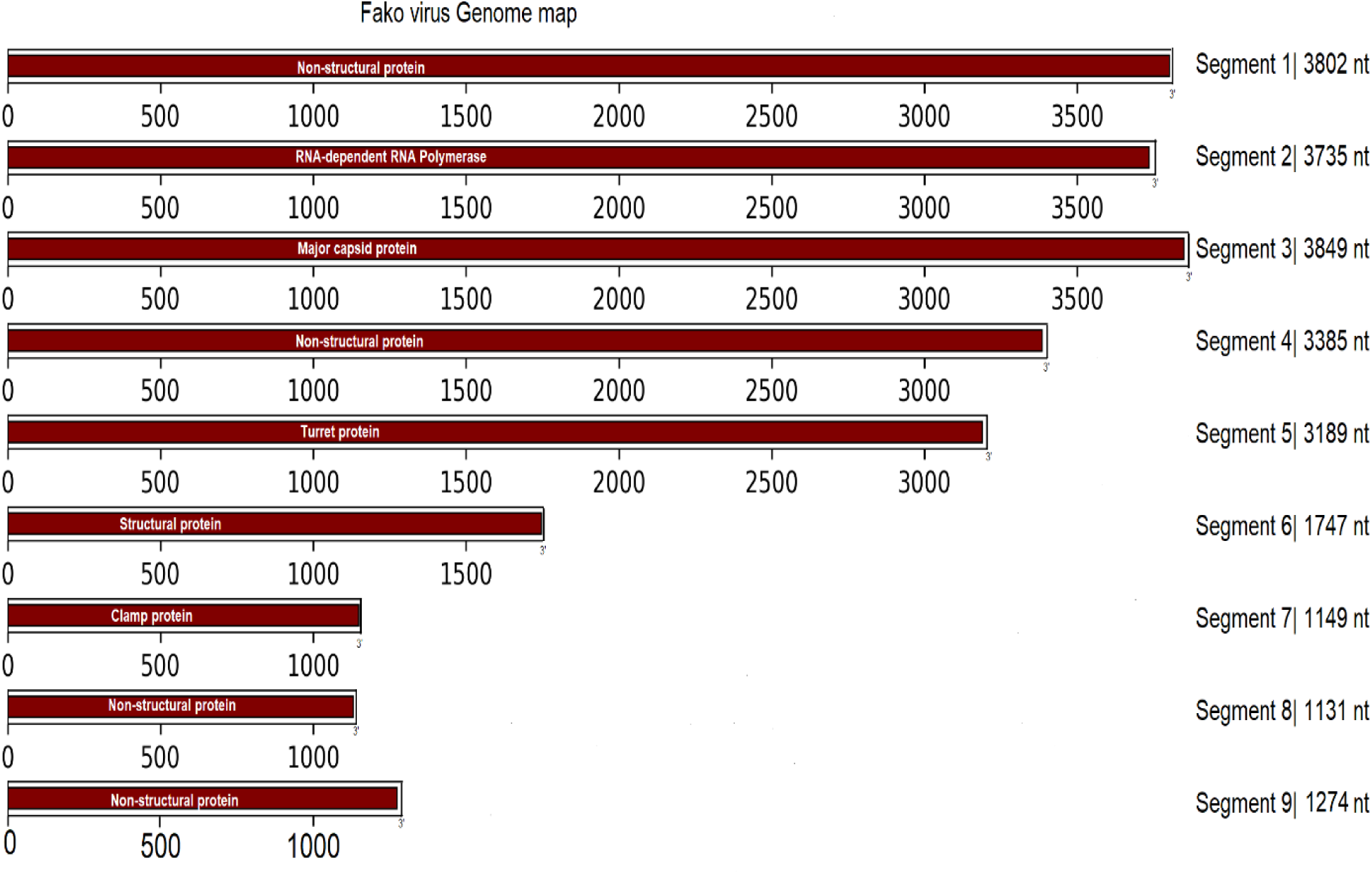
Genome map of Fako virus (FAKV) showing nine segments of varying lengths, with each segment encoding a separate open reading frame.

Phylogenetic analysis of Cell Fusing Agent Virus (CFAV) revealed that the recovered genome clustered with CFAV strains previously reported from Africa, indicating close genetic relatedness across the continent. The branching in the phylogenetic tree was strongly supported, reflecting the conserved nature of this insect-specific flavivirus.

**Figure 10.**
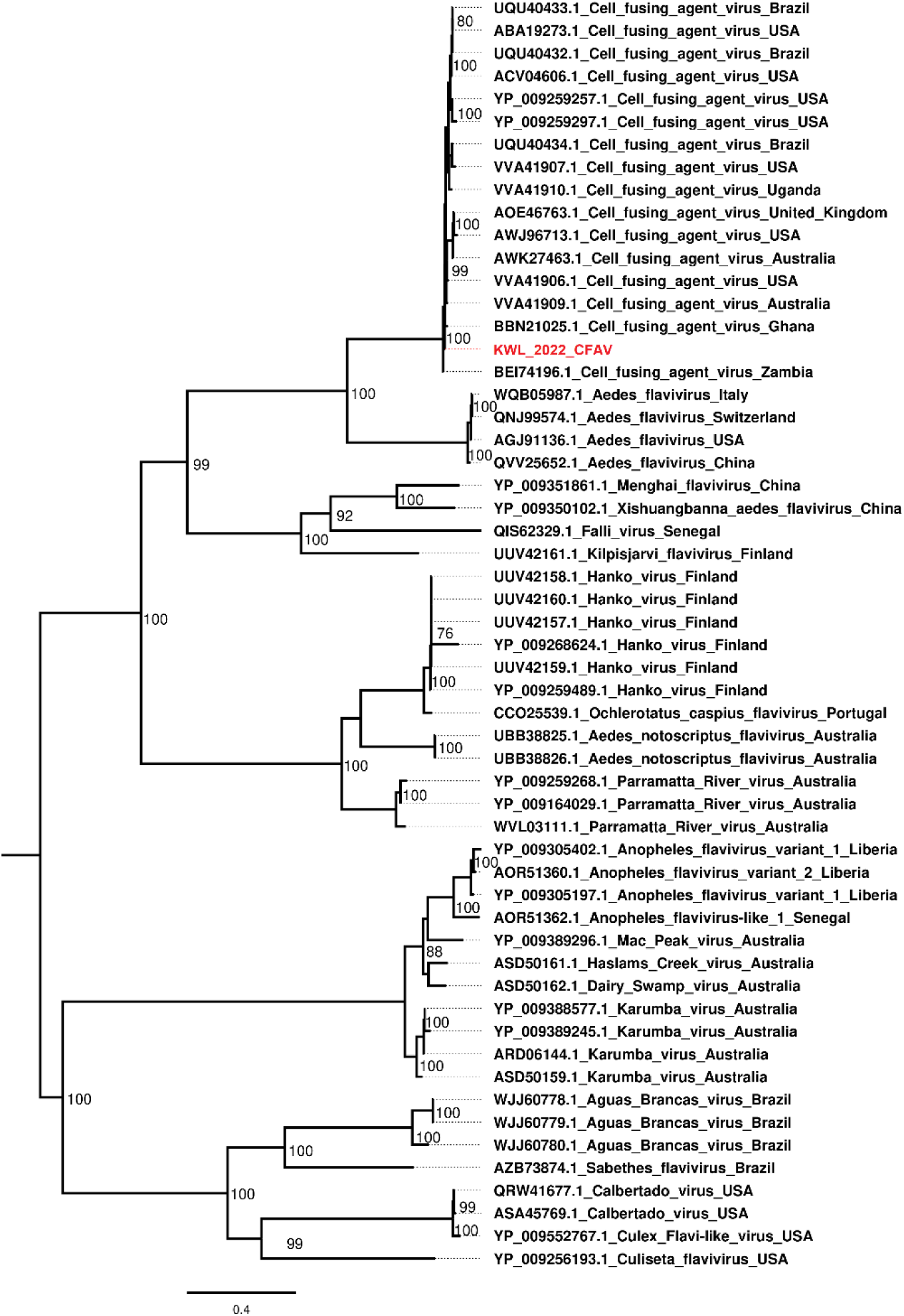
Maximum likelihood phylogenetic tree of Cell Fusing Agent Virus (CFAV). Bootstrap values were calculated from 1000 replicates.

## Discussion

The virome of *Aedes aegypti* is increasingly recognized as being dominated by insect-specific viruses (ISVs), which are unable to infect vertebrates due to their strict host tropism yet may influence mosquito biology and arbovirus transmission dynamics. ISVs are found across multiple viral families, including Flaviviridae, Iflaviridae, Partitiviridae and Reoviridae reflecting both deep evolutionary relationships and potentially distinct ecological roles within mosquito hosts. This study adds to the growing evidence that ISVs constitute a core component of *Ae. aegypti* viromes and highlights the need to understand their broader biological and epidemiological significance.

The detection of Aedes partiti-like virus 1 (AePLV) expands the growing evidence that partitiviruses are stable components of mosquito viromes. Members of the family Partitiviridae are double-stranded RNA viruses with typically bisegmented genomes encoding the RNA-dependent RNA polymerase (RdRp) and capsid protein (Li et al., 2023; Xia et al., 2018). Although historically associated with plants and fungi, partiti-like viruses are increasingly reported in insects, particularly mosquitoes, suggesting long-term host adaptation. Their frequent detection across diverse geographic regions and mosquito populations implies persistent infections, potentially maintained through vertical transmission. While their functional role in mosquitoes remains unclear, recent studies suggest that persistent ISV infections may influence host immunity and vector competence by modulating antiviral pathways.

Formosus virus (FORV) and Tesano Aedes Virus (TeAV) belong to the family Iflaviridae, a group of positive-sense single-stranded RNA viruses widely distributed among insects. Iflaviruses typically establish persistent, often asymptomatic infections and are efficiently maintained in mosquito populations. The repeated detection of TeAV in Aedes aegypti across African regions supports its classification as a mosquito-adapted virus. Importantly, emerging evidence indicates that TeAV may suppress dengue virus replication (Amoa-Bosompem et al., 2020), highlighting a possible indirect role in shaping arbovirus transmission dynamics. Similarly, FORV contributes to the expanding diversity of mosquito-associated iflaviruses, reinforcing the notion that iflaviruses are key constituents of the Aedes core virome with potential ecological and epidemiological relevance.

The identification of Cell fusing agent virus (CFAV), a classical insect-specific flavivirus (cISF), underscores the widespread circulation of insect-restricted flaviviruses in Aedes populations. CFAV is evolutionarily distinct from mosquito-borne pathogenic flaviviruses but shares genomic organization and replication strategies (Chiuya et al., 2021). Experimental studies have demonstrated that it can interfere with the replication of medically important flaviviruses, including dengue and Zika viruses, through mechanisms such as superinfection exclusion and immune priming (Martin et al., 2019). Its presence in natural mosquito populations therefore has important implications for arbovirus ecology, as co-infections with ISFs may influence transmission efficiency and outbreak potential.

The detection of Fako virus (FAKV) places it within the family Reoviridae, a group of segmented double-stranded RNA viruses known to infect a wide range of arthropods. FAKV is characterized by a multi-segmented genome (nine segments), consistent with mosquito-associated reoviruses (Auguste et al., 2014; Reinisch et al., 2000). Reoviruses are thought to establish persistent infections and may be vertically transmitted, facilitating long-term maintenance within vector populations. Although the biological impact of FAKV on mosquito fitness and vector competence remains poorly understood, segmented ISVs such as FAKV contribute substantially to virome complexity and may influence host antiviral responses through continual immune stimulation.

The coexistence of insect-specific viruses from the families Partitiviridae, Iflaviridae, Flaviviridae, and Reoviridae within *Aedes aegypti* underscores the taxonomic breadth and ecological complexity of mosquito viromes. These viruses differ in genome organization and replication strategy but share key features of persistent infection and host restriction. Increasing evidence suggests that such insect-specific viruses are not neutral passengers; rather, they may shape mosquito physiology and vector competence for arboviruses through immune interactions, competitive exclusion, or modulation of cellular environments within the mosquito host. Understanding these interactions is critical for interpreting arbovirus surveillance data and could inform future virome-based strategies for arboviral disease control.

## Conclusion

Collectively, these findings contribute to the growing understanding of mosquito-associated viromes in Africa and emphasize the importance of incorporating insect-specific viruses into arbovirus surveillance and vector biology studies. Characterizing ISVs at the family level provides critical insight into their ecological roles and lays the groundwork for future investigations into their potential application in biological control strategies and arbovirus transmission mitigation.

### Limitations

This study was limited by viral detection being confined to Kwale County, despite processing mosquito samples from Mombasa and Kilifi where no viruses were detected. The use of pooled, cross-sectional metagenomic data limits inference on infection prevalence, viral replication status, and seasonal dynamics. In addition, no arboviruses were detected, and functional interactions between insect-specific viruses and arboviruses could therefore not be assessed in this study.

### Recommendations

- Expand spatial and longitudinal surveillance to capture seasonal and ecological variation in ISV diversity.
- Conduct experimental studies to assess ISV–arbovirus interactions and their effects on vector competence.
- Explore the potential of ISVs for biological control and paratransgenic interventions.

## Ethics approval and consent to participate

Ethical approval was obtained from the Kenya Medical Research Institute (KEMRI) Scientific and Ethics Review Unit (SERU) under protocol number KEMRI/SERU/CCR/4702 and WRAIR# 3101. Permission to conduct the study was granted by the National Council for Science, Technology, and Innovation (NACOSTI).

## Acknowledgements

We thank, Victor Ofula, Dr. Samson Konongoi, Hellen Koka, Dr. Edith Chepkorir, Simon Muhoro and Joseph Katur for their expert contribution in cell culture and data analysis.

## Disclaimer

This Material has been reviewed by the Walter Reed Army Institute of Research. There is no objection to its presentation and/or publication. The opinions or assertions contained herein are the private views of the author, and are not to be construed as official, or as reflecting true views of the Department of the Army or the Department of Defense.

## Availability of data and materials

The sequences of the viruses identified in this study have been submitted to GenBank awaiting accession numbers.

